# Aire-dependent transcripts escape H3K36me3 and Raver2 induced alternative splicing to sustain central immune tolerance

**DOI:** 10.1101/2021.04.08.438874

**Authors:** Francine Padonou, Virginie Gonzalez, Nada Jmari, Julia Maslovskaja, Kai Kisand, Pärt Peterson, Magali Irla, Matthieu Giraud

## Abstract

Aire allows medullary thymic epithelial cells (mTECs) to express and present a large number of self-antigens for central tolerance. Although mTECs express a high diversity of self-antigen splice isoforms, the extent and regulation of alternative splicing events (ASEs) included in their transcripts, notably in those induced by Aire, is unknown. Unexpectedly, and in contrast to Aire-neutral genes, we found that the Aire-sensitive genes exhibit in Aire-positive and negative mTECs, a weak inclusion of ASEs, with about a quarter present in peripheral tissues being excluded from the thymus. We identified Raver2, as a splicing-related factor overrepresented in mTECs and dependent on H3K36me3 marks. We discovered that both Raver2 and methylation of H3K36 promoted ASE inclusion for Aire-neutral genes, leaving Aire-sensitive genes unaffected. Profiling of H3K36me3 revealed its depletion at Aire-sensitive genes, supporting a mechanism, whose setup precedes Aire’s expression and by which Aire-sensitive genes exhibit weak ASE inclusion through the escape of Raver2’s effect. Lack of ASEs in Aire-induced transcripts highlights a role for regulatory T cells in controlling the incomplete Aire-dependent negative selection.

## Introduction

Tolerance against self-tissues is an essential feature of the immune system. It is established in the thymus following the presentation of self-antigen peptides to developing T cells. T cells recognizing their cognate antigen either undergo negative selection by apoptosis, preventing the release of autoreactive T cells, or develop into regulatory T cells (Tregs) able to suppress potential autoreactive T cells in the periphery^1–3^. The subset of medullary thymic epithelial cells characterized by high levels of major histocompatibility complex class II (MHC II) molecules (mTEChi) is essential to the presentation of self-antigens to developing T cells. Indeed, mTEChi have the unique ability to load, onto MHC II molecules, antigenic peptides originated from a wide array of endogenously expressed self-antigens^4,5^, including those controlled by the autoimmune regulator (Aire) and restricted to specific peripheral tissues^4,6^. In mice, invalidation of the *Aire* gene results in a drop of Aire-sensitive gene expression in mTEChi, and the presence of autoantibodies and immune infiltrates directed at multiple peripheral tissues due to impaired negative selection of autoreactive T cells and their harmful activation in the periphery^7^. Mutations in the human *AIRE* gene also cause the multi-organ devastating autoimmune disorder named Autoimmune Polyendocrine Syndrome type 1 (APS1) ^8,9^.

The breadth of the repertoire of self-peptides presented by mTEChi does not only rely on the expression of a high number of self-antigen genes but also on the high diversity of their transcript isoforms whose translation produces multiple protein variants as a result of high rates of alternative splicing^5,10^. Hence, a variety of alternative splicing events (ASEs) are expected to be spliced-in in mTEChi and the resulting processed peptides presented to developing T cells, thereby ensuring the efficient elimination of T cells capable to elicit autoreactive responses against peptides derived from these ASEs in the periphery. Although the expression of a wide range of transcript isoforms has been unambiguously established in mTEChi, the impact and regulation of alternative splicing on the genes controlled by Aire remain to be determined. Notably, it is unknown whether they encode a high transcript-isoform diversity similarly to total mTEChi, or whether the ASEs included in their transcripts equal the diversity of spliced-in ASEs detected for the same genes in their tissues of expression.

We therefore investigated the pattern of ASE inclusion for Aire-sensitive genes in WT and *Aire*-KO mTEChi, as well as in peripheral tissues. Comparison of ASE inclusion for Aire-sensitive versus neutral genes in mTEChi, revealed an unexpected conservation of alternative splicing regulation between Aire-positive and negative mTEC subsets. Differences in the pattern of ASE inclusion with peripheral tissues provided clues on the relative importance of the role of negative selection, in comparison to additional mechanisms, including the generation of suppressive Tregs for the establishment and maintenance of immunological tolerance against Aire-dependent self-antigens. Finally, we identified an epigenetic mechanism that sustains the regulation of alternative splicing in mTECs and explains the patterns of ASE inclusion for Aire-sensitive and neutral genes in these cells.

## Results

### Aire-sensitive genes encode a low diversity of transcript isoforms in mTEChi

To characterize the alternative splicing complexity of Aire-sensitive genes in mTEChi, we determined the diversity of Aire-induced transcript isoforms that result from alternative splicing through the calculation of the median splicing entropy of these genes^11^. For a given gene, splicing entropy was calculated using the levels of transcript-isoform expression obtained by assignment of paired-end sequencing reads to the RefSeq mRNA isoform annotations (**Fig 1A**). The more diverse the transcript isoforms, the greater the associated splicing entropy. Since a sufficient number of mapped junction-spanning reads is necessary for accurate characterization of transcript isoforms, we looked for the minimum expression value below which transcript isoform diversity could not be reliably captured in our mTEChi RNA-seq data. For that purpose, we binned the genes based on expression and calculated their median splicing entropy (**Fig 1B**). We found stable values of splicing entropy for genes with expression levels over 1 FPKM and highly degraded values for genes showing lower expression levels. This prevented the detection of minor transcript isoforms and therefore prompted us to exclude weakly expressed genes for subsequent splicing analyses. Next, we selected the Aire-sensitive genes characterized by a two-fold expression increase in WT versus *Aire*-KO mTEChi (**Fig 1C**) and found that their median splicing entropy was significantly lower than those of Aire-neutral genes and of all genes taken together (**Fig 1D**). This finding thus revealed that Aire-sensitive genes encode a lower diversity of transcript isoforms in mTEChi, denoting lower rates of alternative splicing at these genes.

**Figure 1.**
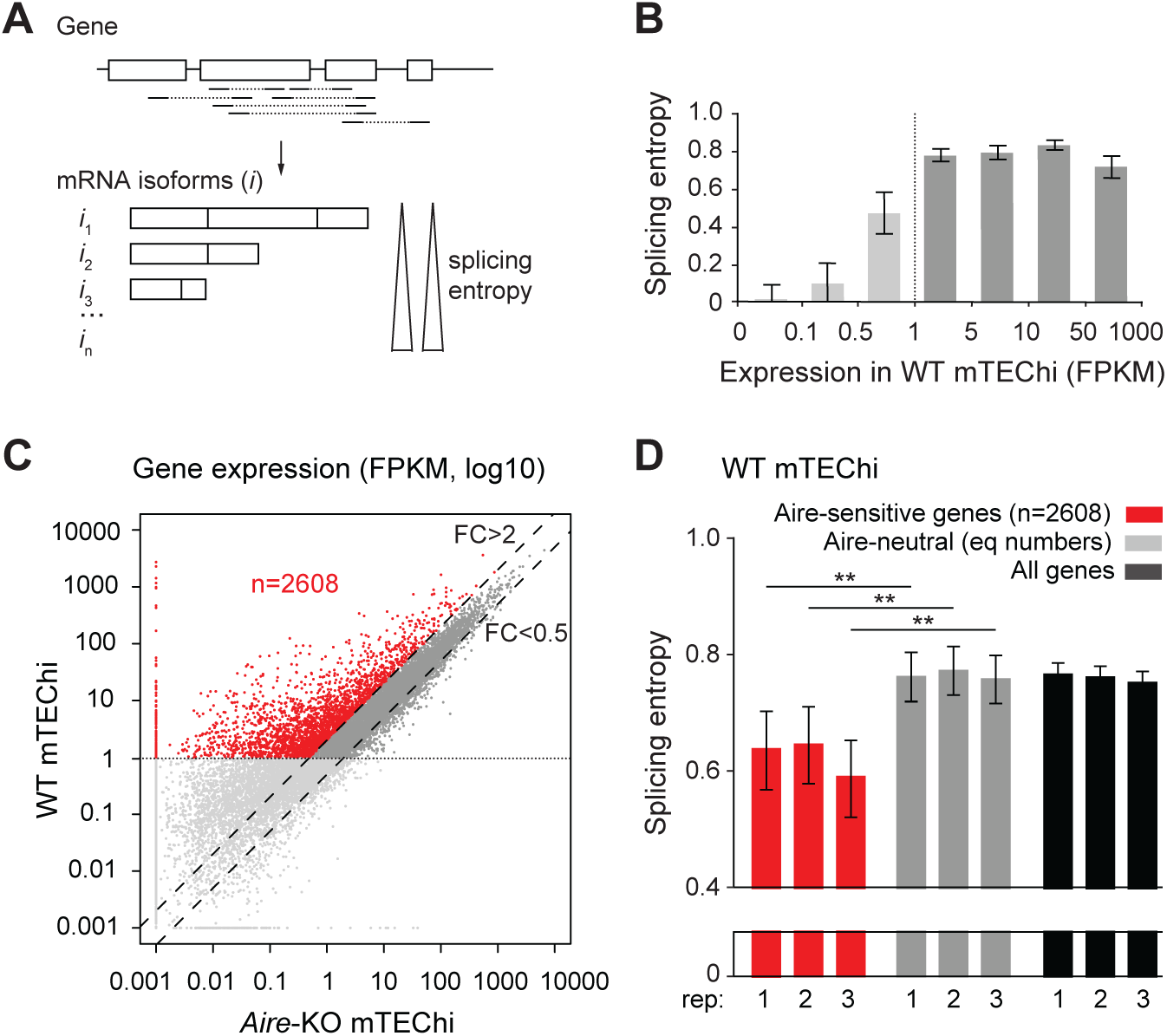
Low splicing entropy of Aire-sensitive genes in mTEChi. **(A)** Schematic representation of a hypothetical gene with mapped paired-end sequencing reads identifying transcript isoforms. The isoform diversity of this gene is evaluated by calculation of its splicing entropy. **(B)** Median splicing entropy of genes binned according to their expression levels. FPKM of 1 corresponds to the threshold over which the transcript isoform diversity can be accurately characterized in our RNA-seq dataset. **(C)** Identification of Aire-sensitive genes upregulated by Aire in WT versus *Aire*-KO mTEChi (FC> 2) and matching the threshold of 1 FPKM in WT mTEChi (red dots, n=2608). Aire-neutral genes (0.5 <FC< 2) with expression levels over 1 FPKM in WT mTEChi are represented by dark gray dots between the dashed lines. **(D)** Median splicing entropy of Aire-sensitive genes, Aire-neutral genes (equal numbers) and all genes in three replicate mTEChi, ** P< 10^−3^ (Wilcoxon test).

### Aire-sensitive genes exhibit weak ASE inclusion in mTEChi

We next sought to determine whether the low diversity of transcript isoforms induced by Aire was sustained by a biased inclusion of ASE. To this end, we first identified all ASEs from the RefSeq mRNA annotation database in parsing its content using rMATS, and then computed their percent splicing inclusion (PSI), i.e., the relative expression of transcript isoforms spliced in versus spliced in or out for each ASE (**Fig 2A**). Comparison of PSI values for Aire-sensitive versus neutral genes in mTEChi revealed a significant imbalance towards lower values (< 0.1), therefore showing that the Aire-sensitive genes exhibit reduced ASE inclusion in mTEChi (Aire-sensitive: PSI< 0.1 (n=163), PSI> 0.1 (n=265); Aire-neutral: PSI< 0.1 (n=93), PSI> 0.1 (n=332); Chisq=3.6×10^−7^) (**Fig 2B**). We then calculated, for Aire-sensitive and neutral genes, the levels of ASE inclusion imbalance, i.e., the number ratio of ASEs showing some level of active inclusion (PSI> 0.1) to ASEs showing no or background inclusion (PSI< 0.1), and found similar reduced levels for Aire-sensitive genes across three replicates (**Fig 2C**, *Left*). Comparison of the levels of ASE inclusion imbalance for Aire-neutral genes between mTEChi and peripheral tissues for which we collected RNA-seq datasets^12^, revealed higher levels in mTEChi, which is in line with reports showing higher rates of alternative splicing in mTEChi^5,10^. In addition, the breadth of the reduction of ASE inclusion between Aire-neutral and Aire-sensitive genes in mTEChi is all the more marked that it is much wider than the reduction observed for Aire-neutral genes between mTEChi and each peripheral tissue (**Fig 2C**, *Right*). Since alternative splicing was reported to be influenced by gene expression in some systems^13^, we calculated the median expression level of neutral genes in each mTEChi replicate and peripheral tissue. We found no significant correlation with the levels of ASE inclusion imbalance, therefore ruling out gene expression as a primary factor responsible for variation of ASE inclusion in our tested samples (**Fig Sup 1**).

**Figure 2.**
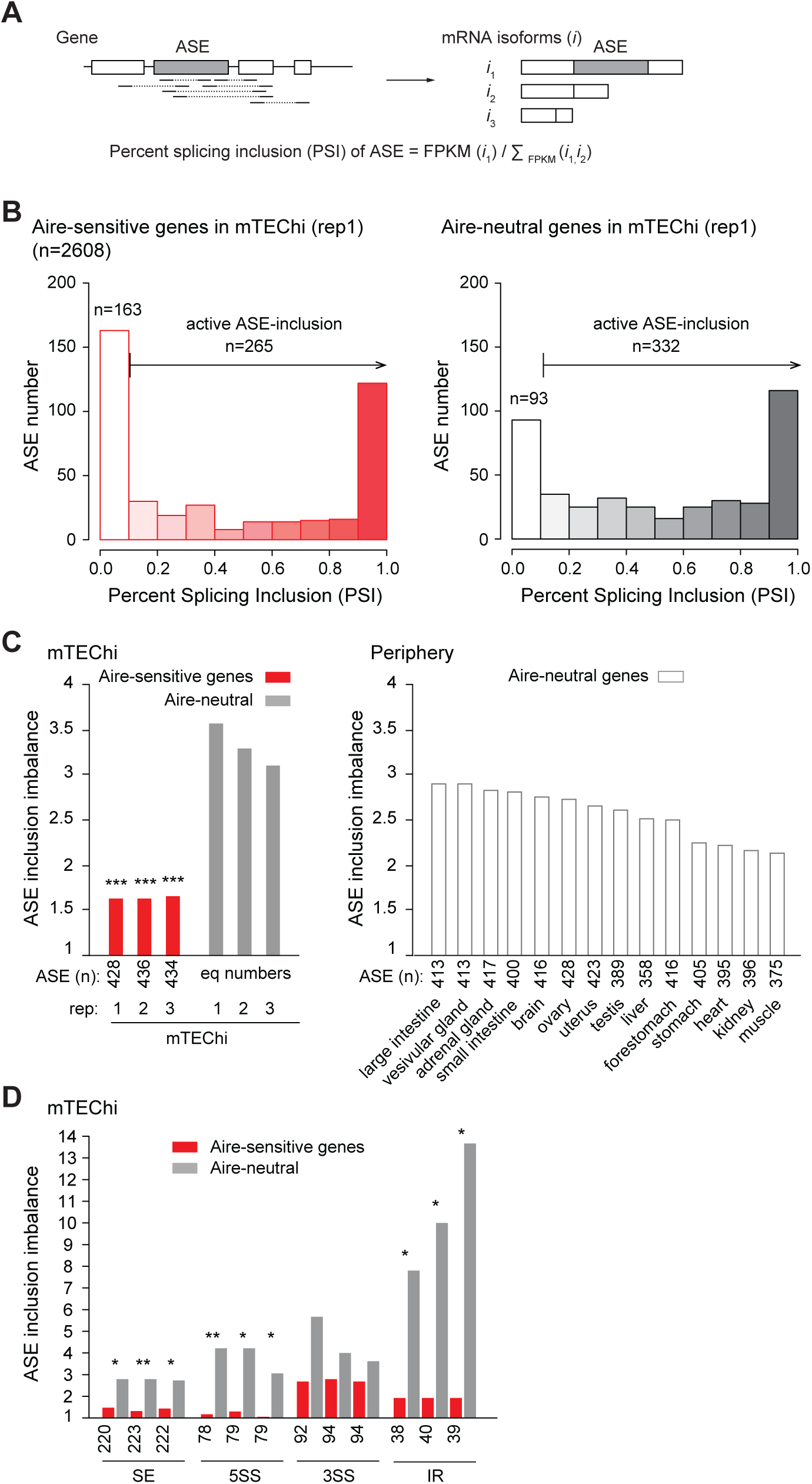
Low levels of ASE inclusion imbalance for Aire-sensitive genes in mTEChi. **(A)** Schematic representation exemplifying the characterization of two transcript isoforms defined by a spliced in (*i*1) and a spliced out (*i*2) ASE, respectively, as well as, of an additional isoform (*i*3) unrelated to the ASE. The PSI value of the ASE is calculated using the specific expression of the transcript isoforms having the ASE spliced in or out. **(B)** Distribution of ASEs according to their PSI values for Aire-sensitive (*Left*) and neutral genes (*Right*) in mTEChi. ASEs with PSI< 0.1 are considered as excluded, whereas ASEs with PSI> 0.1 as present, with some level of active inclusion. **(C)** Levels of ASE inclusion imbalance for Aire-sensitive and neutral genes (equal numbers) in three replicate mTEChi (*Left*) and for Aire-neutral genes in peripheral tissues (*Right*). **(D)** Inclusion imbalance of each type of ASE in three replicate mTEChi. *** P< 10^−4^, **P< 10^−3^, *P< 0.05 (Chi-squared test).

Next, we calculated in mTEChi, the inclusion imbalance of the different types of ASEs, i.e., skipped exon (SE), alternative 5’ splicing site (5SS), alternative 3’ splicing site (3SS) and intron retention (IR) events, and found decreased levels for each type of ASE of Aire-sensitive genes, with SE, 5SS and IR events reaching statistical significance (**Fig 2D**).

Finally, to address whether Aire-sensitive genes also exhibited reduced levels of ASE inclusion in human mTEChi, we isolated mTEChi from human thymic tissues obtained during pediatric cardiac surgery and performed RNA-seq experiments. We identified the minimum expression values over which transcript isoforms could be accurately characterized in these samples, as shown by the example of two individuals (**Fig Sup 2A**). We then identified all ASEs from human RefSeq mRNA annotations and selected human Aire-sensitive and neutral ortholog genes for which we calculated the PSI values of their associated ASEs. Comparison of ASE inclusion imbalance for the Aire-sensitive and neutral genes in one male (indiv 1) and four female (indiv 2 to 5) human individuals showed a marked reduction for Aire-sensitive genes similar to what we found in mice, therefore revealing conservation of ASE inclusion in mTEChi between mice and humans (**Fig Sup 2B**).

Together, these findings showed that Aire-sensitive genes exhibit conserved weak levels of ASE inclusion in mTEChi in contrast to neutral genes that, conversely, exhibit strong levels.

### The weak ASE inclusion of Aire-sensitive genes is a general feature of mTECs

To discriminate whether the reduced inclusion of ASEs of Aire-sensitive genes was directly associated with the Aire’s induction of gene expression or was also observed in the absence of Aire, we selected the genes with two-fold increased expression in WT versus *Aire*-KO mTEChi and with levels of expression over 1 FPKM in both WT and *Aire*-KO mTEChi (**Fig 3A**). We note that meeting the threshold of 1 FPKM in *Aire*-KO mTEChi shrank the selection of Aire-sensitive genes since most of Aire-sensitive genes are inactive or expressed at very low level in the absence of Aire.

**Figure 3.**
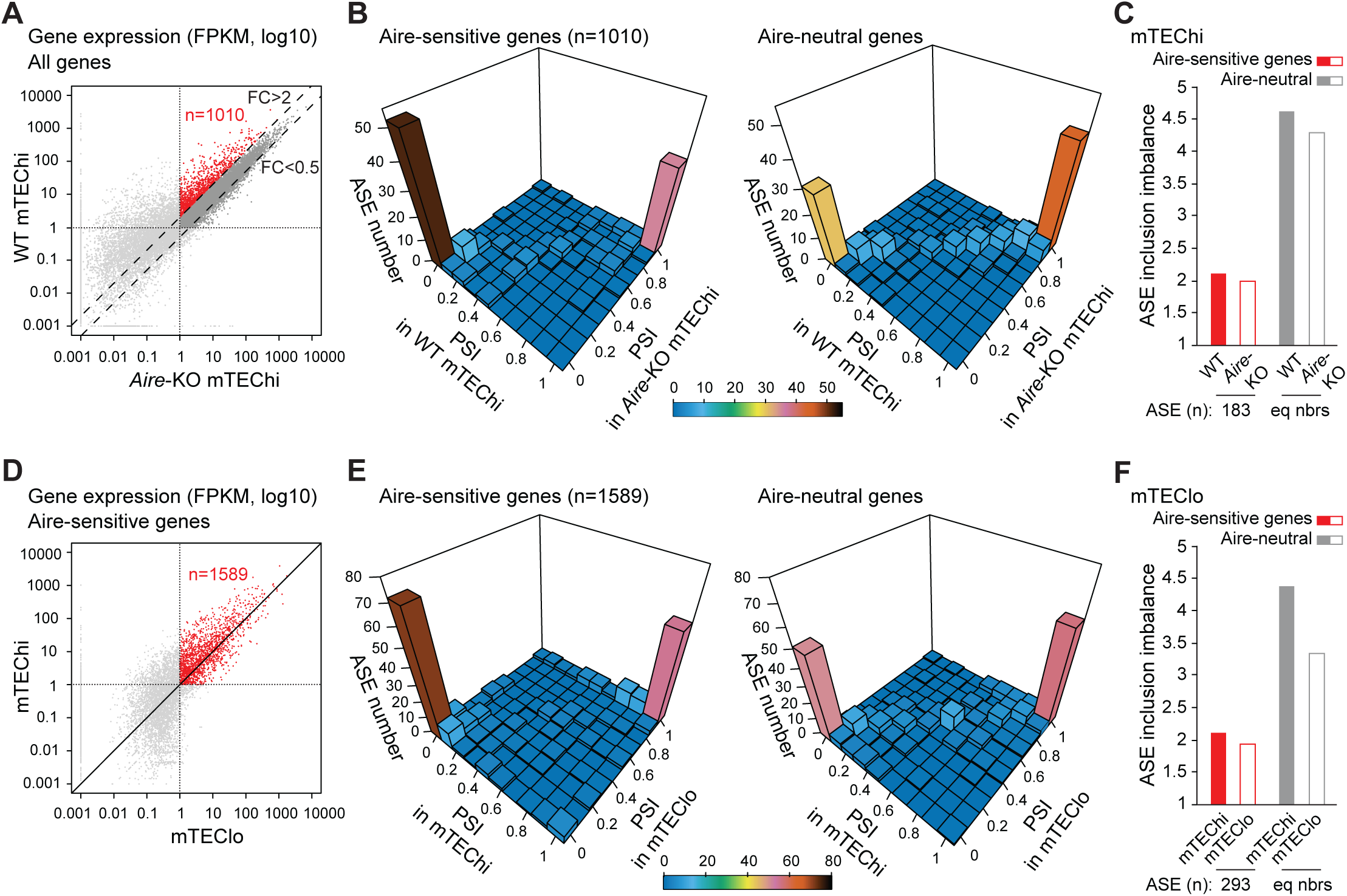
Low levels of ASE inclusion imbalance for Aire-sensitive genes in Aire-negative mTECs. **(A)** Identification of Aire-sensitive genes upregulated by Aire in WT versus *Aire*-KO mTEChi (FC> 2) and matching the threshold of 1 FPKM in WT and *Aire*-KO mTEChi (red dots, n=1010). **(B)** 3D representation of the distribution of ASEs of Aire-sensitive (*Left*) and neutral genes (*Right*) according to their PSI values in WT and *Aire*-KO mTEChi. **(C)** Levels of ASE inclusion imbalance for Aire-sensitive and neutral genes (equal numbers) in WT and *Aire*-KO mTEChi. **(D)** Differential gene expression of Aire-sensitive genes between mTEChi and mTEClo. Red dots show the Aire-sensitive genes with FPKM> 1 in mTEChi and mTEClo (n=1589). **(E)** 3D representation of the distribution of ASEs of Aire-sensitive (*Left*) and neutral genes (*Right*) according to their PSI values in mTEChi and mTEClo. **(F)** Levels of ASE inclusion imbalance for Aire-sensitive and neutral genes (equal numbers) in mTEChi and mTEClo.

We calculated the PSI values of ASEs of the selected Aire-sensitive and neutral genes, and compared their distributions in WT and *Aire*-KO mTEChi using three-dimensional representations (**Fig 3B**). We observed similar PSI values for Aire-sensitive genes in WT and *Aire*-KO mTEChi, as well as for Aire-neutral genes. We then calculated the values of ASE inclusion imbalance and identified similar low levels for Aire-sensitive genes in mTEChi from WT and *Aire*-KO mice, showing that the reduced inclusion of ASEs of Aire-sensitive genes was independent of Aire expression (**Fig 3C**).

Since mTEChi correspond to a stage of differentiation derived from precursor cells in mTEClo that lack Aire expression^14–16^, we asked whether the weak ASE inclusion for Aire-sensitive genes could be a feature already present in mTEClo. To this end, we selected the genes with expression values over 1 FPKM in mTEChi and mTEClo for calculation of PSI values of their associated ASEs (**Fig 3D**). As for the WT versus *Aire*-KO mTEChi comparison, we observed similar PSI values for Aire-sensitive genes in mTEChi and mTEClo, as well as for Aire-neutral genes (**Fig 3E**). Finally, calculation of the values of ASE induction imbalance revealed similar low levels for Aire-sensitive genes in mTEChi and mTEClo (**Fig 3F**), showing that the weak ASE inclusion for Aire-sensitive genes is also a feature of mTEClo.

Together these findings revealed that the weak inclusion of ASEs of Aire-sensitive genes is a general feature of mTECs, conserved upon maturation into mTEChi and independent of Aire expression.

### Aire-sensitive genes show enhanced ASE inclusion in their tissues of expression

Since Aire-sensitive genes are subject to weak ASE inclusion in mTECs, we asked whether they could exhibit a stronger inclusion when their expression is driven by tissue-specific transcriptional mechanisms in the periphery. To address this question, we selected for each tissue in our dataset, the Aire-sensitive genes showing a specific or selective expression by using the SPM (Specificity Measurement) method^17^ as in^18^, and determined the levels of ASE inclusion imbalance. Comparison with mTEChi revealed overall stronger ASE inclusion in peripheral tissues (**Fig 4A**), indicating that ASE inclusion for Aire-sensitive genes is differentially regulated in the periphery. We further identified for each ASE, its PSI values in mTEChi and in the peripheral tissue(s) of specific/selective expression of its corresponding Aire-sensitive gene (**Fig Sup 3A**). We selected the ASEs with PSI values < 0.1 in mTEChi and > 0.1 in the periphery, as exemplified in (**Fig Sup 3A-C**), and found that nearly a quarter of ASEs present in the tissues of specific Aire-sensitive gene expression, were excluded in mTEChi (**Fig 4B**, *Left*). This exclusion of ASEs in mTEChi would therefore increase the risk of release of autoreactive T cells. Conversely, a significantly lower percentage of ASEs present in mTEChi were excluded in the tissues of specific Aire-sensitive gene expression (**Fig 4B**, *Right*). We note that among the latter ASEs, only a minority (about 2 %) showed a full inclusion in mTEChi (PSI> 0.9 in mTEChi and < 0.1 in the periphery) (**Fig Sup 3A**), indicating that only few autoreactive T cells specific for the antigenic epitopes created by the exclusion of these ASEs, would leave the thymus and contribute to the risk for autoimmunity.

**Figure 4.**
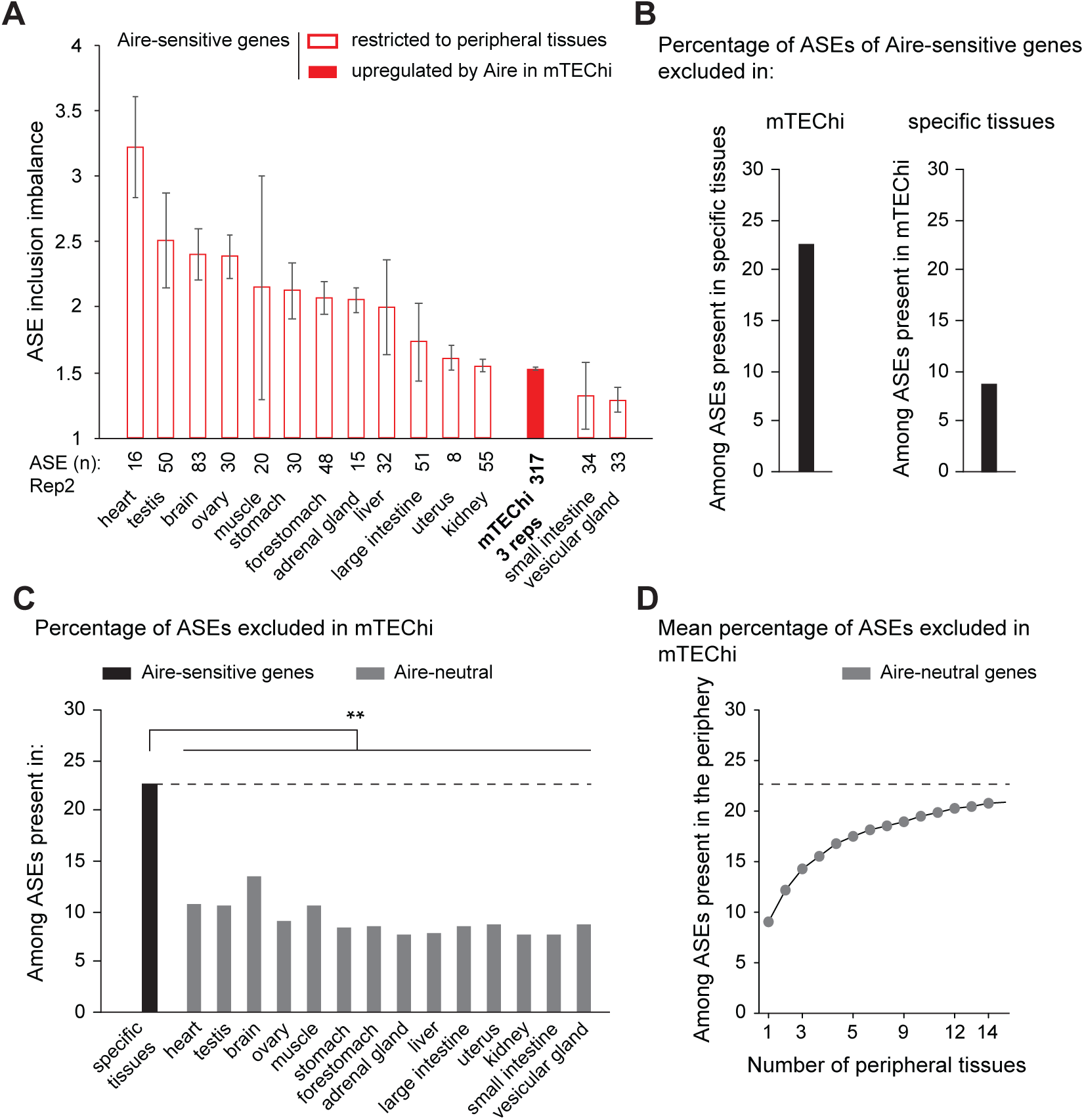
Higher levels of ASE inclusion imbalance for Aire-sensitive genes in their tissues of expression. **(A)** Levels of ASE inclusion imbalance for Aire-sensitive genes in their tissues of expression. Open red bars are for particular peripheral tissues whereas the solid red bar is for mTEChi. **(B)** Percentage of ASEs of Aire-sensitive genes that are excluded in mTEChi among ASEs showing some level of inclusion in the tissues of specific expression (*Left*). The percentage of ASEs excluded in specific tissues among ASEs present in mTEChi is shown (*Right*). **(C)** Percentage of ASE exclusion for Aire-neutral genes in mTEChi among ASEs present in each peripheral tissue or **(D)** in all 1 to 14-permutations of the 14 peripheral tissues in our database, as a mean percentage. The dashed line represents the percentage of ASE exclusion shown in **(B,** *Left*) and **(C)**, **P< 10^−3^.

We next compared for Aire-sensitive and neutral genes the proportion of ASEs excluded in mTEChi among those that are present in peripheral tissues and found significantly lower proportions for neutral genes (**Fig 4C**). Since Aire-neutral genes can be expressed in multiple tissues, increasing their likelihood of undergoing ASE inclusion, we sought to determine for these genes, the proportion of exclusion in mTEChi of their ASEs present in the periphery, considering all tissues together. Hence, we considered all 1 to 14-permutations of the 14 tissues in our dataset, and calculated for each set of permutations of the same number of tissues, the mean proportion of ASEs exclusively excluded in mTEChi (**Fig 4D**). We observed that the proportion of exclusion tended to reach, with the increasing number of considered tissues, the proportion of exclusion detected for Aire-sensitive genes in mTEChi.

Together these findings showed that Aire-sensitive genes exhibit stronger ASE inclusion in their tissues of expression than in mTEChi and that an important proportion of these ASEs present outside the thymus are excluded from mTEChi, similarly to the ASEs of Aire-neutral genes.

### Aire-sensitive genes escape Raver2’s effect promoting ASE inclusion in mTECs

To get insights into the molecular mechanisms that underlie the differential inclusion of ASEs between Aire-sensitive and neutral genes in mTEChi, we searched for the preferential or deprived gene expression of a full set of RNA-binding proteins including factors known to influence RNA splicing, in mTEChi versus peripheral tissues. We identified a handful of RNA-binding proteins showing statistically significant differential expression in mTEChi, including a single factor reported to modulate alternative splicing, namely Raver2^19^ (**Fig 5A** and **Fig Sup 4**). We further showed that Raver2 expression is over-represented in mTEChi and mTEClo, supporting an important role in mTECs (**Fig 5B**).

**Figure 5.**
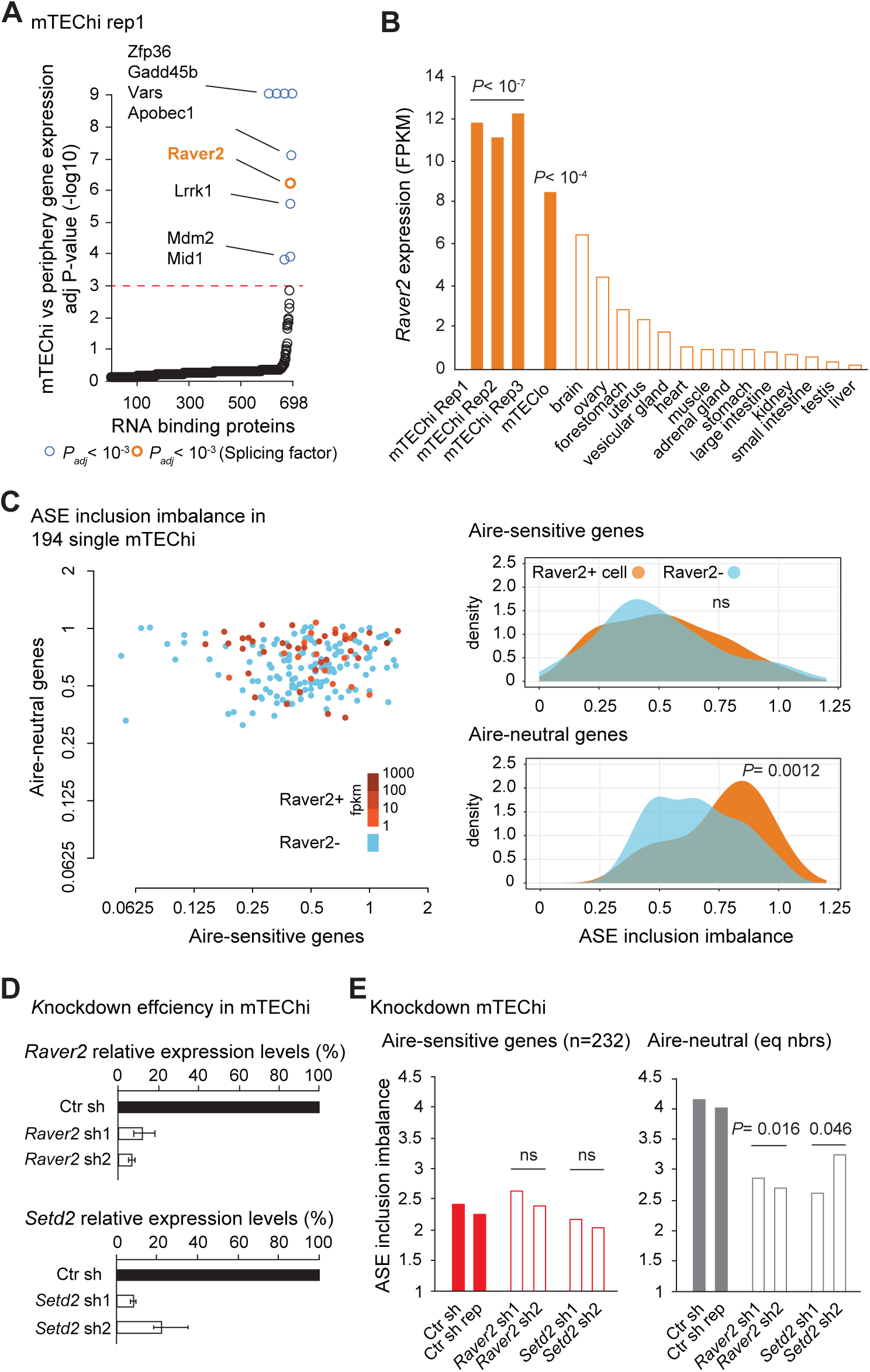
Effect of Raver2 and Setd2 on the level of ASE inclusion imbalance for Aire-sensitive and neutral genes in mTECs. **(A)** Differential expression of genes coding for RNA-binding proteins in mTEChi vs peripheral tissues. The red dashed line represents the threshold for statistical significance (Benjamini-Hochberg adjusted *P*< 0.001). RNA-binding proteins showing significant differential gene expression are represented by colored circles, in orange if the RNA-binding proteins have been reported to be involved in splicing, in blue otherwise. **(B)** *Raver2* expression levels in mTEChi, mTEClo and across peripheral tissues (open bars). Significance of Raver2 over-representation in mTEChi and in mTEClo versus the peripheral tissues is shown. **(C)** Levels of ASE inclusion imbalance (*Left*) and their distributions (*Right*) are shown for Aire-neutral and Aire-sensitive genes in 194 single mTEChi. Blue is for cells negative for Raver2 whereas orange shades are for Raver2-positive cells. Statistical significance assessed by a Student’s test. **(D)** Relative expression levels of *Raver2* (*Top*) and *Setd2* (*Bottom*) in mTEChi infected by lentiviruses containing *Raver2*, *Setd2* or the Ctr (LacZ) shRNAs. **(E)** Levels of ASE inclusion for Aire-sensitive and neutral genes (equal numbers) in mTEChi infected by lentiviruses targeting *Raver2*, *Setd2* (open bars) or the Ctr (LacZ) (solid bars). Statistical significance of cumulative sh1 and sh2 assessed by a Chi-squared test.

To determine whether Raver2 was able to exert an effect on Aire-sensitive or neutral genes, we first sought to correlate its expression to variation of ASE inclusion imbalance across individual mTECs. To this end, we collected full-length single-cell RNA-seq data enabling splicing analyses, that were generated in mTEChi^20^. We calculated, for each cell, the PSI values of ASEs of Aire-sensitive and neutral genes, as well as their levels of ASE inclusion imbalance. Note that the comparison of the distribution of the levels of ASE inclusion imbalance across the individual mTEChi and between Aire-sensitive and neutral genes showed significantly lower levels for Aire-sensitive genes (**Fig Sup 5A**), therefore confirming, at the single-cell level, the difference that we identified at the cell population level in (**Fig 2C** and **Fig Sup 2B**). The absence of Aire effect on the low inclusion of ASEs of Aire-sensitive genes was confirmed by the lack of significant correlation between the levels of ASE inclusion imbalance and those of *Aire* expression across the individual mTEChi (**Fig Sup 5B**). We then identified which cells expressed *Raver2* (threshold> 1 FPKM) (**Fig 5C**, *Left*), and compared the distribution of the levels of ASE inclusion imbalance between the Raver2-positive and negative cells. In contrast to Aire-sensitive genes, we observed a significant association between the presence of Raver2 and the high levels of ASE inclusion imbalance for Aire-neutral genes, strongly suggesting that Raver2 promotes the inclusion of ASEs specifically in Aire-neutral genes (**Fig 5C**, *Right*).

Then, to evaluate the specific impact of Raver2 on ASE inclusion in mTECs, we seeded mTECs onto a 3D organotypic culture system adapted from^21^ and performed infection with concentrated lentiviruses encoding shRNAs targeting *Raver2* (shRNA 1 or 2) or the control LacZ, cloned into a vector expressing GFP as a marker of transduction (**Fig Sup 6**). Three days later, we cell-sorted GFP+ mTECs and quantified *Raver2* expression level. We found a strong knockdown efficiency for each of the two *Raver2* shRNAs with 10 % to 20 % remaining expression in comparison to control knockdown samples (**Fig 5D**, *Top*). To ensure that the newly transcribed isoforms, resulting from potential alternative splicing alterations promoted by *Raver2* knockdown, stabilize and accumulate to be accurately detected, we isolated GFP+ mTECs five days after infection. RNA-seq experiments on these cells revealed a significant reduction of the levels of ASE inclusion imbalance for Aire-neutral genes (**Fig 5E**, *Right*), whereas no effect was detected on Aire-sensitive genes (**Fig 5E**, *Left*). This finding demonstrated that Raver2 promotes the inclusion of ASEs of Aire-neutral genes and that the Aire-sensitive genes escape its effect.

Raver2 is a heterogeneous nuclear ribonucleoprotein (Hnrnp) that has been reported to bind to the polypyrimidine track-binding protein (Ptb) ^22,23^ which regulates alternative splicing at regions enriched in H3K36me3 through interaction with H3K36me3 adaptor proteins^24,25^. To evaluate the effect of H3K36 methylation on ASE inclusion in mTECs, we performed knockdown of Setd2 that is recognized as the only enzyme able to methylate H3K36 into H3K36me3 in somatic cells^26^. We first validated the efficient knockdown of two shRNAs targeting *Setd2* (**Fig 5D**, *Bottom*) and then performed RNA-seq experiments to determine the levels of ASE inclusion imbalance in the knockdown samples. As for *Raver2*, *Setd2* knockdown resulted in a significant reduction of ASE inclusion for Aire-neutral genes whereas no effect was detected for Aire-sensitive genes (**Fig 5E**).

Together these findings showed that Aire-sensitive genes escape Raver2’s effect promoting ASE inclusion in mTECs, and suggested that the effect on Aire-neutral genes may require H3K36me3 as an anchor for this factor.

### Aire-sensitive genes are devoid of H3K36me3 and H3K36me3-associated ASE inclusion in mTEClo

Since the weak ASE inclusion for Aire-sensitive genes is already present in mTEClo (**Fig 3D-F**), we sought to determine whether a potential escape of Raver2’s effect due to a lack of H3K36me3 could apply for such a mechanism in these cells. We looked for H3K36me3 enrichment by Chip-seq experiments, and found very low levels of H3K36me3 at Aire-sensitive genes in comparison to all genes (**Fig 6A** and **Fig Sup 7A**). We then selected the genes with expression values over 1 FPKM and the presence of H3K36me3 marks within the last third of their gene-bodies, and compared the levels of ASE inclusion imbalance to those of the counterpart genes lacking H3K36me3 (**Fig 6B**). We observed a significant increase of ASE inclusion for genes harboring H3K36me3 marks, very likely reflecting the functional link that we detected between increased H3K36me3 and stronger ASE inclusion in mTECs (**Fig 5E**, *Right*). Thus, these findings supported the assumption that the weak ASE inclusion observed for Aire-sensitive genes in mTEClo was due, at least in part, to the lack of H3K36me3 deposition and the subsequent impairment of Raver2 recruitment.

**Figure 6.**
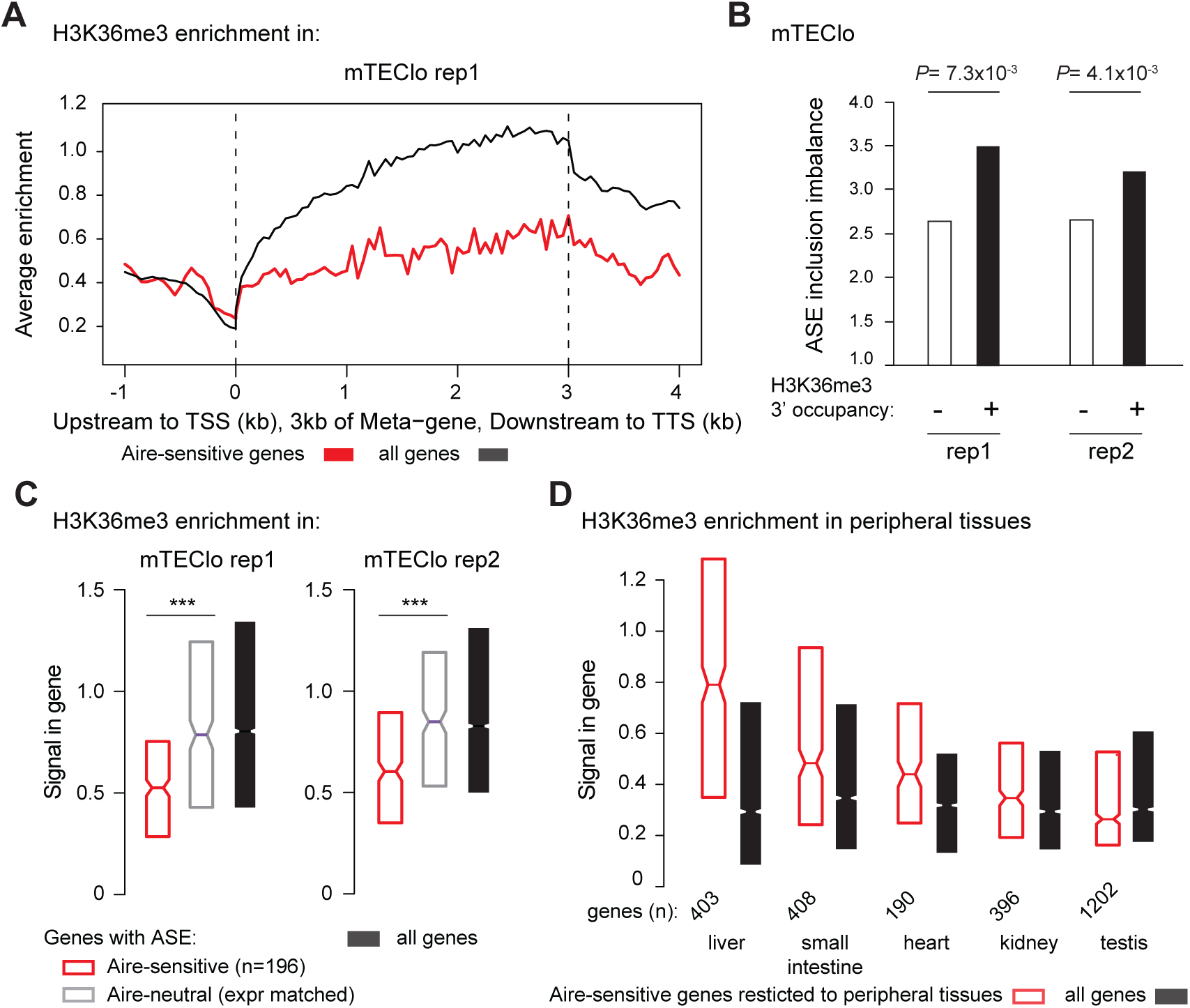
Aire-sensitive genes show low levels of H3K36me3 correlated with low levels of ASE inclusion imbalance in mTEClo. **(A)** Metagene profiles of the average normalized enrichment of H3K36me3 for Aire-sensitive genes (red) and all genes (black) in mTEClo. **(B)** Levels of ASE inclusion imbalance for genes with and without H3K36me3 in the last third of their gene-bodies, in two replicate mTEClo. **(C)** Median enrichment of H3K36me3 for Aire-sensitive genes and neutral genes (expression matched), as well as all genes in two replicate mTEClo, *** *P*< 10^−13^ (Wilcoxon test). **(D)** Median enrichment of H3K36me3 for Aire-sensitive genes in their tissues of expression and for all genes.

However, since H3K36me3 is involved in transcription^27,28^, we asked whether the lack of H3K36me3 at Aire-sensitive genes was specific to these genes or associated with their low expression. To address this question, we selected a set of Aire-neutral genes with expression levels matching those of Aire-sensitive genes, and measured their enrichment for H3K36me3. We found significantly lower levels of H3K36me3 at Aire-sensitive genes, therefore showing that the lack of H3K36me3 was specific to these genes and unrelated to their low expression (**Fig 6C**). In addition, a comparison of the levels of ASE inclusion imbalance for the same Aire-sensitive and expression-matched neutral genes showed a clear reduction of the former (**Fig Sup 7B**), confirming that the weak ASE inclusion for Aire-sensitive genes is not a reflection of their low expression levels.

Finally, to address whether the lack of H3K36me3 deposition at Aire-sensitive genes was a differential feature between mTEClo and peripheral tissues, we collected H3K36me3 Chip-seq data generated from mouse tissues, as part of the ENCODE project^29^. For each tissue, we measured the enrichment of H3K36me3 at all genes and found that Aire-sensitive genes with a specific or selective expression showed similar to higher levels of H3K36me3 in comparison to all genes (**Fig 6D** and **Fig Sup 7C**). This finding is consistent with the observation that Aire-sensitive genes exhibit enhanced ASE inclusion, globally, in the periphery, but showed a poor correlation with ASE inclusion in each individual tested tissue (**Fig 4A**), indicating that H3K36me3 may be permissive to tissue-specific mechanisms of alternative splicing without orchestrating a coordinated tissue-independent pattern of the latter.

## Discussion

In our present study, we demonstrated that, while mTEChi express a wide range of self-antigen splice isoforms, Aire-upregulated transcripts exhibit a low diversity and a weak inclusion of ASEs. We showed that this low ASE inclusion was independent of Aire’s action on gene expression and resulted from a mechanism that was sustained by the specific depletion of H3K36me3. The linked mechanism between the decrease of H3K36me3 and the reduction of ASE inclusion that we propose, was consistent with the report showing that alternative exons, subjected to exclusion or partial inclusion, have lower H3K36me3 signals than constitutive exons in *Caenorhabditis elegans*^30^. In addition, a recent report showed that high inclusion levels of skipped exon (SE) ASEs correlate with the strong enrichment of H3K36me3 in the exon flanking regions in mice^31^.

Unlike Aire-sensitive genes, we showed that the genes neutral to the effect of Aire in mTECs were characterized by an enrichment of H3K36me3 and a strong enhancement of ASE inclusion. Identification of Raver2 as a splicing-related factor overrepresented in mTECs and able to promote ASE inclusion led us to propose a mechanism by which ASE inclusion for Aire-neutral genes is driven by the coordinated action of Raver2 and the splicing factor Ptb that is specifically recruited at H3K36me3-rich regions. This mechanism was supported by the observation that Ptb is not only able to repress alternatively spliced exons^32^ but also enhances the inclusion of a large number of ASEs, as shown by the effect of *Ptb* knockdown on ASE inclusion at the genomic scale^24^. In addition, Raver2 was characterized as a potent modulator of the splicing activity of Ptb^19^, therefore opening the possibility that Raver2 may function through interaction with Ptb to enhance ASE inclusion for genes with H3K36me3 enrichment. The role of Raver2 in mTECs is further strengthened by the higher expression of *Ptb* in mTEChi (105 FPKM) and mTEClo (91 FPKM) than in peripheral tissues (median expression: 31 FPKM).

Hence, through depletion of H3K36me3, the self-antigen genes controlled by Aire likely escape the effect of Raver2/Ptb, and thereby generate transcripts with fewer ASEs. This weak ASE inclusion indicated that the Aire-induced transcripts do not include all the ASEs that are spliced-in in the periphery. Lack of presentation of Aire-dependent antigenic peptides corresponding to ASEs absent from mTEChi, is likely to result in a number of autoreactive T cells leaving the thymus and potentially able to elicit autoimmune reactions. Hence, the negative selection triggered by Aire is not as comprehensive as originally thought, therefore highlighting a role for complementary peripheral tolerance mechanisms, notably mediated by Tregs. Tregs selected against peptides derived from constitutive exons of Aire-sensitive genes in mTEChi, would then suppress the autoimmune responses triggered against immunogenic ASE-derived peptides. The underlined role of Tregs in Aire-mediated tolerance is in line with previous studies showing the importance of Aire-induced self-antigen expression in the selection and development of suppressive Tregs^33–35^. In humans, the study of APS1 patients revealed a role for AIRE in the development of Tregs^36,37^, as well as in the enforcement of immune tolerance by directing differentiation of autoreactive T cells into the Treg lineage^38^. This latter mechanism would thus provide the possibility for autoreactive T cells directed against immunogenic ASE-derived peptides, to convert into Tregs, therefore preventing potential autoimmune reactions and favoring Treg-associated suppressive responses.

Our study also revealed that the proportion of ASEs selectively excluded in mTEChi is similar between Aire-sensitive and neutral genes. This indicated that the lack of H3K36me3 at Aire-sensitive genes, and therefore the absence of Raver2 recruitment, would result in a certain degree of autoreactivity that Tregs may control similarly to autoreactive reactions against immunogenic peptides derived from ASEs of Aire-neutral genes. Hence, the weak ASE inclusion for Aire-sensitive genes in mTECs may be sufficient to impose, with the support of Tregs, immune tolerance against the full set of Aire-dependent self-antigens. If Aire-mediated immunological tolerance was entirely under the control of the sole mTEChi-dependent negative selection, which happens not to be the case, the full set of ASEs of Aire-sensitive genes would be present in mTEChi with the full diversity of tissue-specific splicing mechanisms activated. Such a broaden activation would very likely conflict with the proper function and physiology of mTEChi and represent a tremendous energetic cost, most probably unreachable.

## Materials and Methods

### Mice

*Aire*-deficient (C57BL/6) mice, previously obtained by D. Mathis and C. Benoist (Harvard Medical School, Boston, MA), were bred and maintained at Cochin Institute. Wild-type B6 mice were purchased from Charles River laboratory. Animal housing and experiments were conducted in specific-pathogen-free conditions according to the protocols and guidelines of the French Veterinary Department under procedures approved by the Paris-Descartes Ethical Committee for Animal Experimentation (decision CEEA34.MG.021.11).

### Isolation of medullary thymic epithelial cells

Thymi of 4-6 weeks old mice were trimmed of fat, cut into pieces and digested by mechanic coercion to release thymocytes. Enzymatic digestion was first performed with collagenase D (1 mg/mL) (Roche) and DNase I (1 mg/mL) (Sigma) for 30 min at 37 °C, then with Collagenase/Dispase (2 mg/mL) (Roche) and DNase I (2 mg/mL) at 37 °C up to the obtention a single-cell fraction. The cells were then filtered through a 70 μm cell strainer and resuspended in PBS containing 1% FBS and 5 mM of EDTA. mTECs were then isolated from the obtained cell fraction using the following methods:

#### Isolation of mTEChi and mTEClo

Thymic stromal cells were enriched by CD45 depletion of thymocytes using magnetic CD45 MicroBeads (Miltenyi Biotec) and the magnetic cell sorter (autoMACS, Miltenyi Biotec). The depleted cell fraction was then stained for 20 min at 4 °C with a cocktail of fluorophore-labeled antibodies CD45-PerCPCy5.5 (1:50) (Biolegend), Ly51-PE (1:800) (Biolegend) and I-A/E-APC (MHCII) (1:1,200) (eBioscience) for sorting of mTEChi/lo (CD45^-^Ly51^-^I-A/E^high/low^) on a FACSAria III instrument (BD Bioscience), as previously described^18,39^.

#### Isolation of total mTECs for 3D organotypic culture

As above, a CD45 depletion of thymocytes using magnetic beads was first performed. The depleted cell fraction was then stained for 20 min at 4 °C with the CD45-PerCPCy5.5 antibody (1:50) (Biolegend) and the EpCAM-FITC antibody (G8.8 clone) (1:500) (Biolegend) for FACS sorting of CD45^-^EpCAM^+^ cells. The sorted fraction was sequentially stained for 20 min at 4 °C with the Ly51-PE antibody (1:800) (Biolegend) and depleted of Ly51-positive cells using anti-PE Microbeads (Miltenyi Biotec), then stained with the I-A/E-APC antibody (MHCII) (1:1,200) (eBioscience) and positively selected for I-A/E-positive cells using anti-APC Microbeads (Biolegend). The obtained final cell fraction consisted of total mTECs (CD45^-^EpCAM^+^Ly51^-^I-A/E^+^).

### shRNA-containing lentivirus production

pLKO.1 plasmids bearing the shRNA sequences (sh1: TRCN0000181363 and sh2: TRCN0000242035 for *Raver2*; sh1: TRCN0000238533 and sh2: TRCN0000238536 for *Setd2*; Ctr sh: TRCN0000072240 for LacZ) (Sigma) were subcloned into the lentiviral pLKO.3G vector containing an eGFP cassette (Addgene #14748), by transferring the BamHI-NdeI restriction fragments containing the shRNAs. Lentiviruses were produced into HEK293T cells by using the calcium phosphate co-transfection method for each specific subcloned lentiviral pLKO.3G vector with the packaging plasmids gag/pol (Addgene #14887) and VSV-G (Addgene #14888). HEK293T were grown in Dulbecco’s modified Eagles medium (DMEM) with high glucose (4500 mg/L), supplemented with 10% fetal bovine serum (FBS), pen/strep antibiotics. HEK293T cells were transfected at 70-80% confluence in two T175 flasks. The transfection solution was prepared with 500 µl of 1 M CaCl2 (Sigma), the lentiviral vector (32 µg), the packaging plasmids (VSV-G: 16 µg, gag/pol: 24 µg) and DNAse-free water (Invitrogen) for a final volume of 2 mL. Then, 2 mL of HEPES-buffered saline pH 7.0 (2X for transfection) (VWR) was added dropwise to the previous mixture, under constant agitation. The obtained solution was kept at room temperature for 15 minutes and subsequently equally transferred to the HEK293T flasks. The next day, the medium was changed. 48 h after transfection, the culture medium containing the lentiviral particles was collected and ultracentrifuged at 25,000 rpm for 90 min at 4 °C using a Beckman SW28 ultracentrifuge rotor (Beckman Coulter) for the obtention of concentrated virus, that was resuspended in cold PBS 1X. Viral titration was performed in HEK293 cells. For individual evaluation of shRNAs, mTECs were transduced at a multiplicity of infection of 10.

### 3D organotypic culture of primary mTECs

#### Preparation of the fibrin gel

The 3D organotypic co-culture system described in^21^ was set up using a ∼1.2-mm-thick viscose coated nonwoven fibrous fabric: Jettex 2005/45 (Orsa) that was placed as a scaffold into 12-well filter inserts (polyester capillary membrane with 3 μm pores) (Dutscher). A semi-solid fibrin gel inoculated in the insert, that serves as support for the culture system was prepared seeding human dermal fibroblasts (200,000) re-suspended in a fibrin gel made of fibrinogen (Merck-Millipore), thrombin (Merck-Millipore) and aprotinin (Euromedex) which prevents precocious fibrinolysis by serine proteases secreted by fibroblasts. Human dermal fibroblasts were generated from explant cultures of de-epidermized dermis as described^40^ and grown up to passage 4 in DMEM/F-12 (Thermofisher) complemented with 10% FBS and pen/strep antibiotics. The fibrin gel polymerized at 37 °C for one hour or two, forming a smooth upper surface for the culture of primary mTECs. A pre-culture of five days is required for the fibroblast activation and is set up by the submersion of the polymerized gel in medium 1 containing DMEM/F-12, 10 % FBS, pen/strep antibiotics, L-ascorbic acid (50 μg/mL) (Sigma) and TGF-β (1 ng/mL) (R&D Systems). The medium 1 was changed every day. On the day of mTEC seeding, medium 1 was replaced by medium 2 containing DMEM high glucose + DMEM/F12 1:1 (v/v) with 10 % FBS, pen/strep antibiotics, cholera toxin (10^−10^ M) (Sigma), hydrocortisone (0.4 μg/μL) (Sigma), L-ascorbic acid (50 μg/mL), aprotinin (500 U/mL) and RankL (80 ng/mL) (R&D Systems).

#### mTEC seeding and lentiviral infection

Freshly isolated mTECs were seeded at the top of the fibroblast-containing fibrin gel with the addition of 10 µl of concentrated shRNA-bearing lentiviruses that we produced against *Raver2*, *Setd2* and the Ctr (LacZ). The next day, medium 2 was replaced by medium 3 that is similar to medium 2 but containing reduced amount of aprotinin (250 U/mL) and increased amount of RankL (100 ng/mL). The RankL supplemented in mediums 2 and 3 helped maintain mTEChi maturity.

#### Isolation of knockdown mTECs

Three days after infection, a first part of the 3D organotypic co-culture was terminated by cutting out half of the fibrin gel to evaluate the *Raver2* and *Setd2* knockdown efficiencies by RT-qPCR. Two days later, the other half of the fibrin gel was processed to evaluate the impact of *Raver2* and *Setd2* knockdown on ASE inclusion of Aire-sensitive and neutral genes by RNA-seq. Using small dissecting scissors, the fibrin gel portions were cut into small pieces to be properly digested in an enzymatic solution of Collagenase/Dispase (0.5 mg/mL) (Roche) and DNase I (0.5 mg/mL) at 37 °C for 15 min for the obtention a single-cell fraction. After three rounds of centrifugation and clean-up in PBS containing 1% FBS and 5 mM of EDTA, the single-cell fraction was stained for 20 min at 4 °C with the I-A/E-APC (MHCII) antibody (1:1,200) (eBioscience) and for 5 min at 4 °C with the DAPI viability dye solution (1 µg/mL) (Sigma) to identify dead cells. We then sorted by flow cytometry DAPI^-^I-A/E^+^GFP^+^ cells corresponding to viable lentivirus-infected mTECs.

### shRNA knockdown efficiency

RNA extraction from knockdown mTECs cultured in the 3D organotypic system, was performed using the Single Cell RNA Purification Kit (Norgen Biotek). First-strand cDNA was synthesized using SuperScript IV VILO Master Mix (Thermofisher). cDNA was used for subsequent PCR amplification using the viia7 Real-time PCR system (Thermofisher) and the Fast SYBR Green Master Mix (Thermofisher). Knockdown efficiency was assessed by comparing the level of expression of *Raver2* (forward primer: AACCAGAAGACACCGCAGAG; reverse: TCTCCCAAGAGTGAAGTCTGATT) and *Setd2* (forward primer: CAGCATGCAGATGTAGAAGTCA; reverse: TCCAGGACAAAGGTGTTCG) between the knockdown and control samples using *Gapdh* (forward primer: GGCAAATTCAACGGCACAGT; reverse: AGATGGTGATGGGCTTCCC) for normalization.

### RNA-seq of medullary epithelial cells

Total RNA of knockdown mTECs cultured in the 3D organotypic system, was extracted using the Single Cell RNA Purification Kit (Norgen Biotek). Total RNA of mTEChi isolated from WT or *Aire*-KO mice, was extracted with TRIzol (ThermoFisher) and used to generate the third replicate of two independent paired-end RNA-seq datasets that we previously generated and deposited in the GEO database (GSE140683). We used mTEClo paired-end RNA-seq data that we previously generated (GSE140815). Single-cell full-length and paired-end (2×125bp) RNA-seq data of mTEChi, were obtained from the GEO database (GSE114713). For mTECs in the 3D organotypic system, polyA-selected transcriptome libraries were constructed using the SMART-seq v4 Ultra Low Input RNA Kit (Takara) combined to the Nextera XT DNA Library Preparation Kit (Illumina) and sequenced on the Illumina NextSeq 500 machine as paired-end data (2×150bp). Sequences were deposited in the GEO database as GSEXX. For WT and Aire-KO mTEChi, polyA-selected transcriptome libraries were constructed using the TruSeq Stranded mRNA Library Prep (Illumina) and sequenced on the Illumina HiSeq 2000 machine as paired-end data (2×100 bp). Sequences were deposited in the GEO database as GSEXX. All paired-end reads of the different datasets were homogenized to 2×100 bp by read-trimming and mapped to the mm10 genome using the TopHat 2 program^41^ with default parameters. Levels of gene and transcript isoform expression were determined using the cufflinks 2 program^42^ that assembles all reads on the RefSeq transcript annotations and estimates their abundances. We also used cuffdiff 2^43^ to compute differential gene expression between WT and *Aire*-KO mTEChi and between mTEChi and mTEClo.

### Human medullary epithelial cell isolation and RNA-seq

The thymi from cardiac surgery pediatric patients (the ethical permission #170/T-I from Ethics Review Committee on Human Research of the University of Tartu) were minced in 1xPBS to release the thymocytes. Then treated four times with collagenase/dispase (1:200) and DNaseI (1:1000) digestion, and the cell fractions were further dissociated with gentleMACS treatment. All fractions were combined and filtered through the 100μm Falcon cell strainer. The cells were collected by centrifugation, washed, and transferred to Optiprep density gradient. The upper cell layer was combined into one tube, washed, and counted resulting in 0.4-6.5x 10^8^ thymic cells. The cell fraction was then depleted with human CD45 beads (100ul beads per 2×108 cells; Miltenyi Biotec) using autoMACS. After the CD45 depletion, the cells were sorted with BD FACSAria for mTEC MHC^hi^ (CD45^-^ CDR2^-^ EpCAM^+^ HLA-DR^hi^) directly into TRIzol (ThermoFisher) solution, RNA was isolated with miRNAeasy Mini Kit (Qiagen) with DNase treatment. The RNA quality was assessed with TapeStation (Agilent D1000) before the processing with SMART-Seq v4 Ultra Low Input RNA Kit for Sequencing, and the library preparation with Illumina Nextera XT DNA Library Preparation Kit. The paired-end sequencing (2×150bp) was performed on the Illumina NextSeq500. Sequences were deposited in the GEO database as GSEXX. The fastq files were mapped to the hg19 genome with TopHat 2. Levels of gene and transcript isoform expression were determined using cufflinks 2 and expression data as well as GTF files were used for ASE analysis.

### H3K36me3 Chip-seq experiment and analysis

Nano-ChIP-seq on H3K36me3 was performed as previously described^44^ on 50,000 isolated mTEClo. ChIP-seq libraries were prepared with the TruSeq ChIP Sample Preparation Kit (Illumina) and 2×75 bp paired-end reads were sequenced on an Illumina HiSeq 2000 machine. Sequences were deposited in the GEO database as GSEXX. Alignment to the mm10 genome was done using Bowtie 2^45^. Duplicate alignments were removed using Samtools^46^ and the command line: “samtools view -S -hf 0×2 alignments.sam | grep -v “XS:i:” | foo.py > filtered.alignments.sam” the foo.py script been available online: “https://www.biostars.org/p/95929/”. Peak calling for a sample (in comparison to the inputs) was done using MACS2 callpeak^47^ and the options -f BAMPE, --SPMR and –broad. Enrichment to the inputs, was calculated using MACS2 bdgcmp and the option -m FE. For multi-tissue comparison, H3K36me3 Chip-seq data generated by the Bing Ren’s lab (http://renlab.sdsc.edu/), as part of the ENCODE project^29^, were obtained from the GEO database: GSE31039 for heart, kidney, liver, small intestine and testis of adult (8-week old) mice. These data were available as signal (bigWig) and peak (bed) files processed on the mm9 genome following ENCODE standards. Finally, we used the CEAS distribution^48^ on mTEClo and tissue data to calculate the average H3K36me3 enrichment within gene bodies and their upstream and downstream regions.

### Splicing entropy calculation

The splicing entropy of a specific gene is a measure of diversity of the transcript isoforms that result from alternative splicing for this gene^11^. The higher the number of transcript isoforms, the higher the splicing entropy. For a particular gene, the expression values of its transcript isoforms, computed by cufflinks, are used to calculate the splicing entropy:

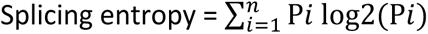

n is the number of isoforms and P*_i_* is the proportion that each isoform (*i*) contributes to the overall expression of the gene. We calculated the splicing entropy of each gene in our samples of interest and used the median splicing entropy for sample-to-sample comparison.

### ASE inclusion analysis

We used the rMATS program (Multivariate Analysis of Transcript Splicing) ^49^ to identify all ASEs from the mm10 or hg19 RefSeq mRNA annotations database, including skipped exon (SE), alternative 5’ splicing site (5SS), alternative 3’ splicing site (3SS) or intron retention (IR). For a particular ASE, we computed the percent splicing inclusion (PSI), i.e., the relative expression of transcript isoforms spliced in versus spliced in or out, using the R-script that we developed (GitXXX):

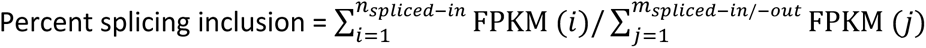

n_spliced-in_ is the number of isoforms with the ASE spliced in. m_spliced-in/-out_ is the number of isoforms with the ASE spliced in or spliced out. To compare the PSI values between different set of genes and/or samples, we determined the splicing inclusion imbalance by dividing the number of ASEs showing some level of active inclusion (PSI> 0.1) by the number of excluded ASEs, i.e., showing no or background inclusion (PSI< 0.1).

### Multi-tissue comparison analysis

We considered 14 homogeneously sequenced RNA-seq datasets (paired-end) of mouse peripheral tissues (Brain, Liver, Kidney, Adrenal Gland, Heart, Ovary, Testis, Stomach, Forestomach, Small intestine, Large Intestine, Muscle, Uterus, Vesicular Gland) generated by^12^, and obtained from the NCBI BioProject database (PRJNA375882). The reads were mapped to the mouse reference genome (mm10) with TopHat 2, using default parameters. Expression levels of transcript isoforms were calculated using Cufflinks. We determined the tissue specificity (one tissue of restricted expression) or selectivity (two-to-three tissues of restricted expression) of Aire-sensitive genes, by using the specificity measurement (SPM) and the contribution measurement (CTM) methods^17^, as described in^18^. For each considered gene, the SPM and CTM values were evaluated based on the level of gene expression in each tissue. A gene was considered tissue-specific for a particular tissue if its SPM value in the tissue was > 0.9. A gene was considered tissue-selective (2 or 3 tissue), if its SPM values were > 0.3 in those tissues and its CTM value for the corresponding tissue > 0.9. If the previous conditions were not met, the gene was left unassigned.

## Acknowledgments

We thank Drs. D. Mathis and C. Benoist (Harvard Medical School) for *Aire*-KO (B6) mice. This work was supported by the Agence Nationale de la Recherche (ANR) 2011-CHEX-001-R12004KK to M.G., IHU-Cesti funded by the «Investissements d’Avenir» ANR-10-IBHU-005 as well as by Nantes Metropole and Region Pays de la Loire to M.G., and EJP-Rare Disease JTC2019 program TARID project funded by the ANR (ANR-19-RAR4-0011-5) to M.G., by Estonian Ministry of Social Affairs to P.P.. The work was also supported by the University of Tartu Center of Translational Genomics (SP1GVARENG) and the Estonian Research Council grant PRG377 to P.P. and the Marie Curie Actions (Career Integration Grants, CIG_SIGnEPI4Tol_618541) to M.I.. We would like to thank the members of the ‘Genomic’IC’ core facility head by Franck Letourneur at Cochin Institute, Paris, France, for RNA-seq data production, as well as Ms. Maire Pihlap for her excellent technical assistance in preparing sequencing samples, Dr Mario Saare for his help in analyzing human mTEC data and Nicolas Richard for his help with the single-cell analysis. F.P. was supported by the Labex IGO (project «Investissements d’Avenir» ANR-11-LABX-0016-01) and the RFI Bioregate grant (ThymIPS) from la Region Pays de la Loire to M.G.. F.P. and M.G. designed the study and wrote the manuscript; F.P., V.G. and N.J. performed most of the experimental work; J.M. and K.K. performed experiments. F.P. and M.G. performed bioinformatics analyses; P.P. and M.I. provided key material, datasets and edited the manuscript.

## Figures Sup legends

**Figure Sup 1.**
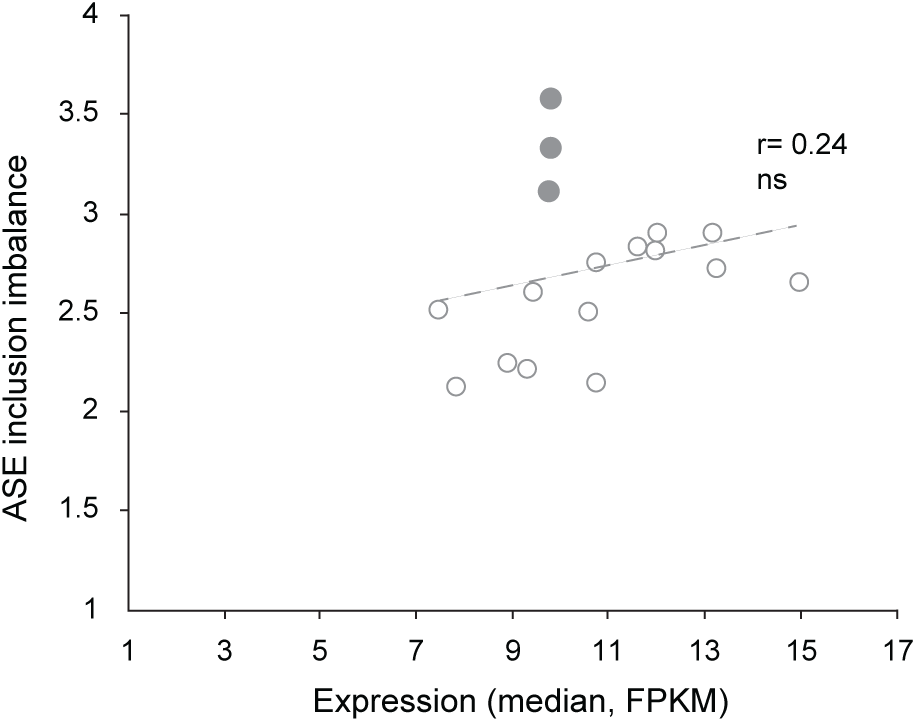
Lack of correlation between the levels of ASE inclusion imbalance and of expression of Aire-neutral genes across tissue and mTEChi samples. Levels of ASE inclusion imbalance for Aire-neutral genes relative to their median gene expression in peripheral tissues (opened circles) and three replicate mTEChi (closed circles). The dashed line stands for the linear regression (not significant) across all samples. Pearson correlation.

**Figure Sup 2.**
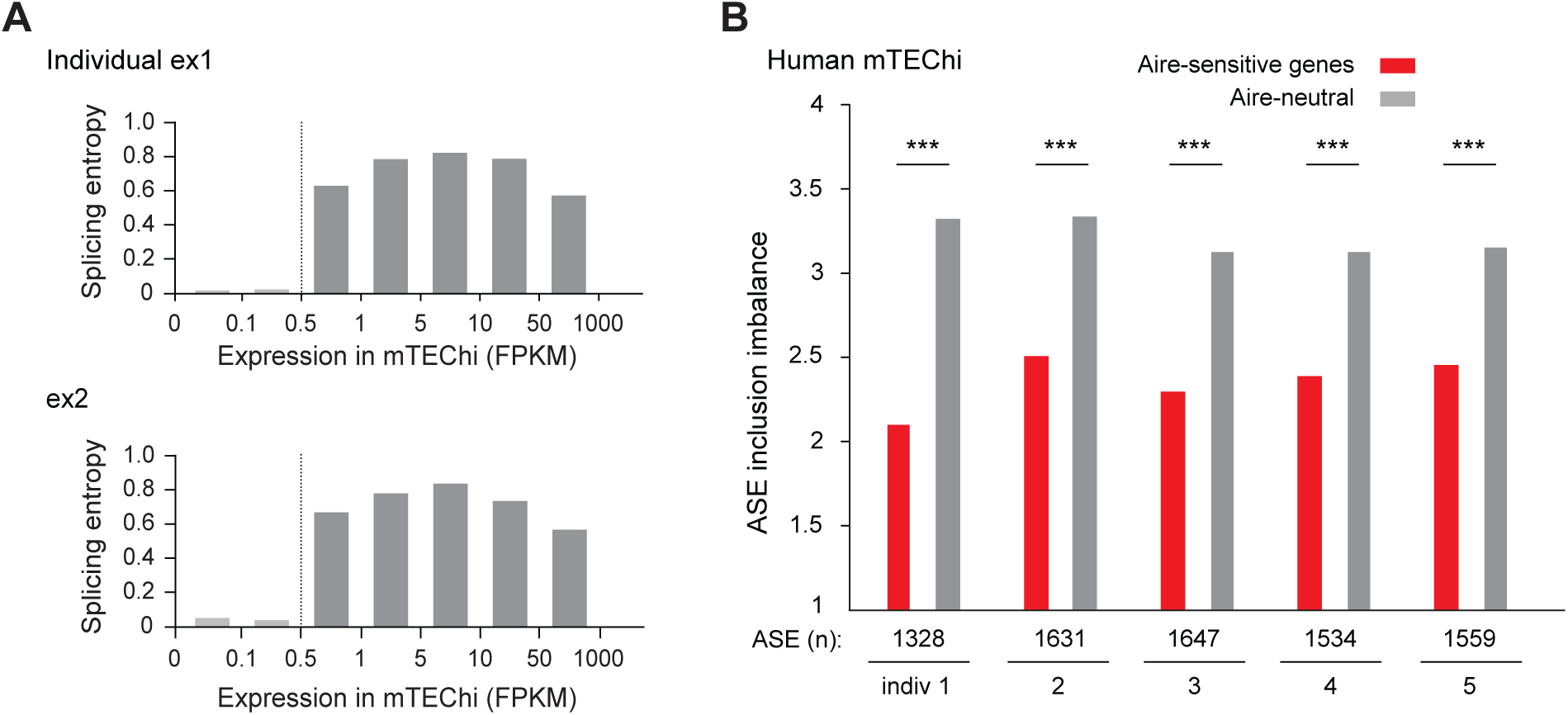
Low levels of ASE inclusion imbalance for Aire-sensitive genes in human mTEChi. **(A)** Median splicing entropy of genes binned according to their expression values. FPKM of 0.5 corresponds to the threshold over which the transcript isoform diversity can be accurately characterized in our RNA-seq dataset. Example of two individuals is shown. **(B)** Levels of ASE inclusion imbalance for Aire-sensitive and neutral genes in mTEChi of five different individuals, *** P< 10^−4^ (Chi-squared test).

**Figure Sup 3.**
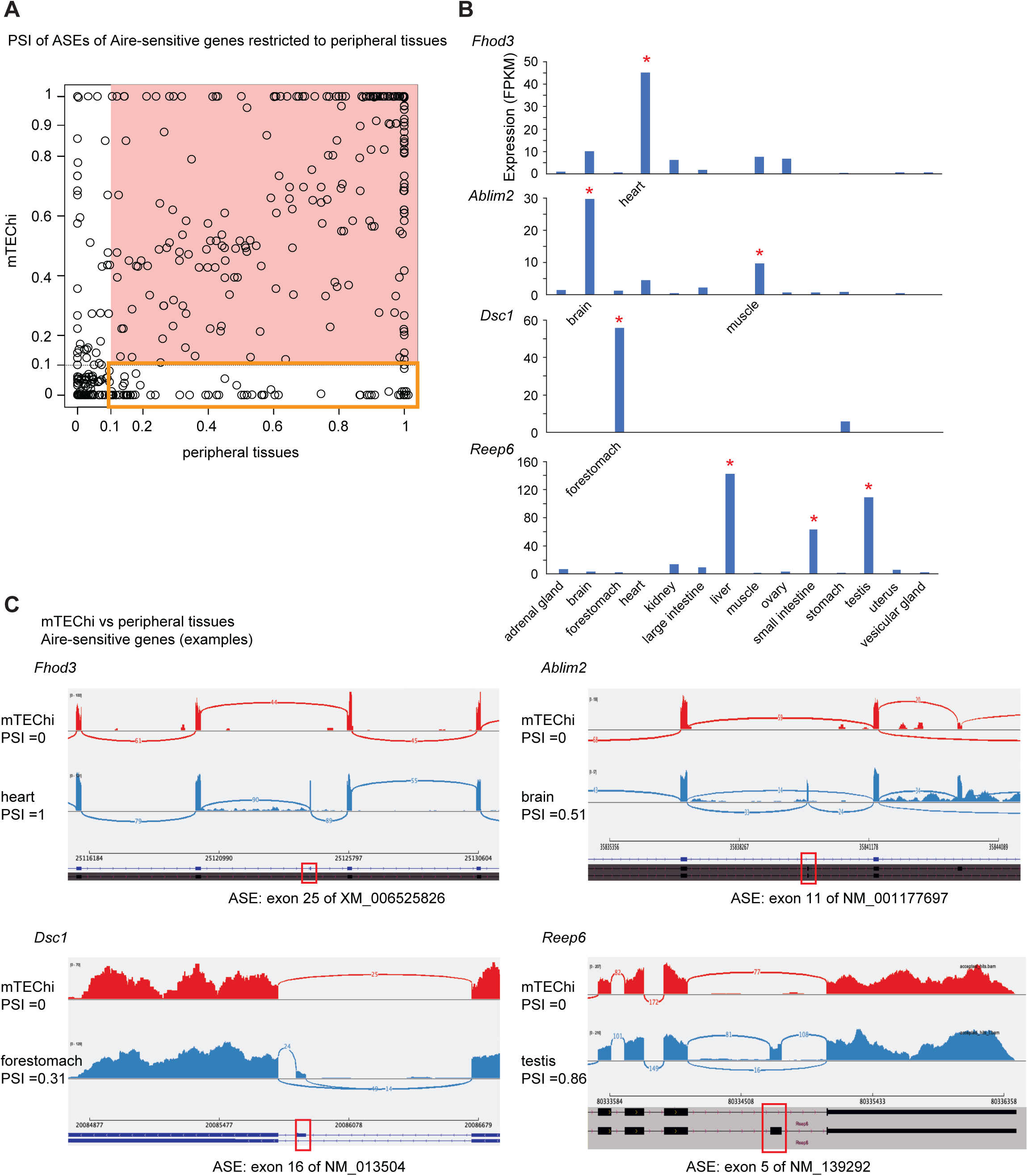
PSI values of ASEs of Aire-sensitive genes in mTEChi and in their tissues of expression. **(A)** Each ASE is represented by a circle. ASEs excluded in mTEChi or in the periphery (PSI< 0.1) are shown on a white background, otherwise on a salmon-colored background. The ASEs present in the periphery (PSI> 0.1) and excluded in mTEChi (PSI< 0.1) are framed by an orange square. **(B)** Level of expression of *Fhod3*, *Ablim2*, *Dsc1* and *Reep6* (taken as examples of Aire-sensitive genes with a specific or selective peripheral expression) from RNA-seq data of mTEChi and 14 mouse tissues. *Fhod3* is specific to the heart, *Dsc1* to the forestomach; *Ablim2* is selective to the brain and muscle, *Reep6* to the liver, small intestine and testis. **(C)** Shashimi plots of the above genes show ASE exclusion (PSI< 0.1) in mTEChi (red) and some level of inclusion (PSI> 0.1) in the tissues of expression (blue). Arcs representing splice junctions connect exons.

**Figure Sup 4.**
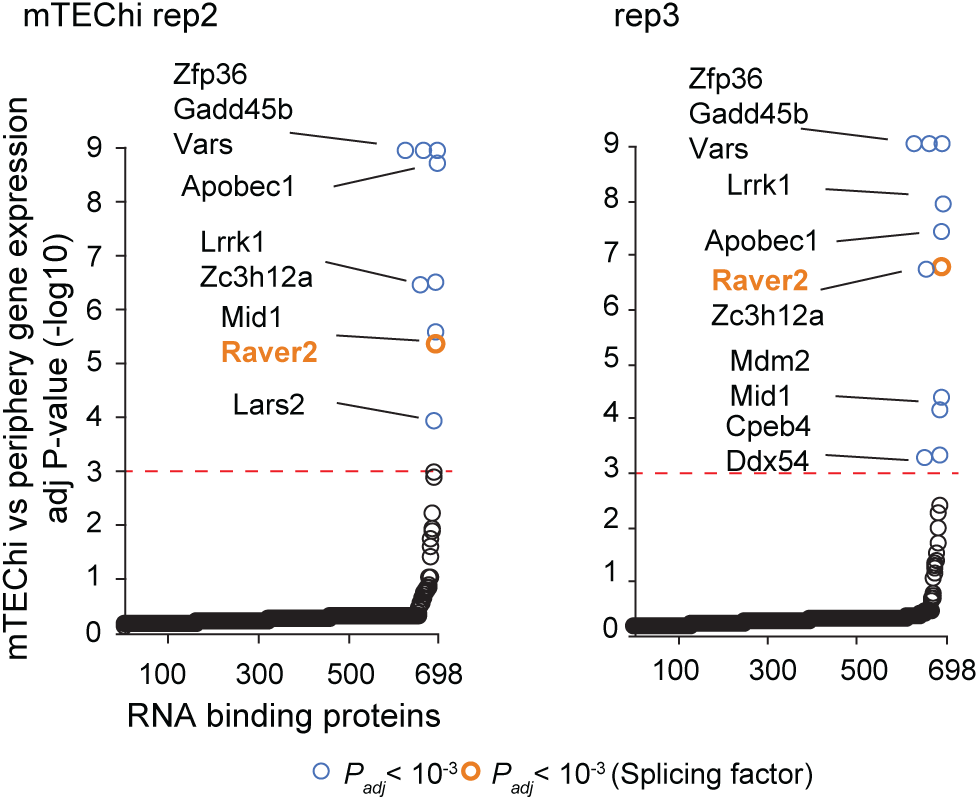
Differential expression of genes coding for RNA-binding proteins in two replicate mTEChi vs peripheral tissues. The red dashed line represents the threshold for statistical significance (Benjamini-Hochberg adjusted *P*< 0.001). RNA-binding proteins showing significant differential gene expression are represented by colored circles, in orange if the RNA-binding proteins have been reported to be involved in splicing, in blue otherwise.

**Figure Sup 5.**
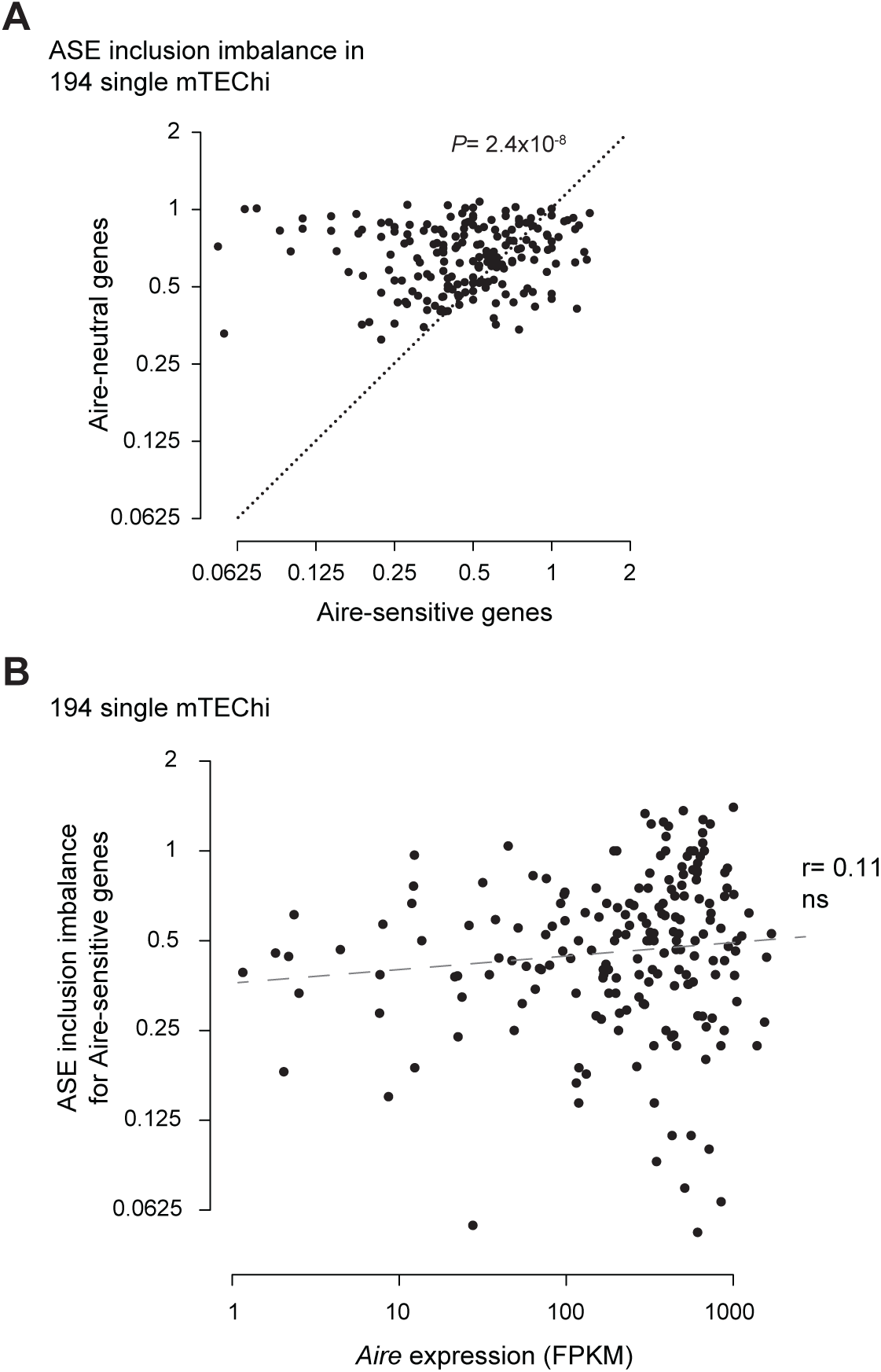
Low levels of ASE inclusion imbalance for Aire-sensitive genes in single mTEChi. **(A)** Levels of ASE inclusion imbalance are shown for Aire-neutral and Aire-sensitive genes in 194 single mTEChi that are positive for Aire (FRKM> 1). The dotted diagonal line represents similar ASE inclusion for the two gene sets. The distribution of cells is significantly skewed towards lower levels of ASE inclusion imbalance for Aire-sensitive genes. Student’s t-test. **(B)** Levels of ASE inclusion imbalance for Aire-sensitive genes relative to their median gene expression in single mTEChi. The dashed line stands for the linear regression (not significant) across all samples. Pearson correlation.

**Figure Sup 6.**
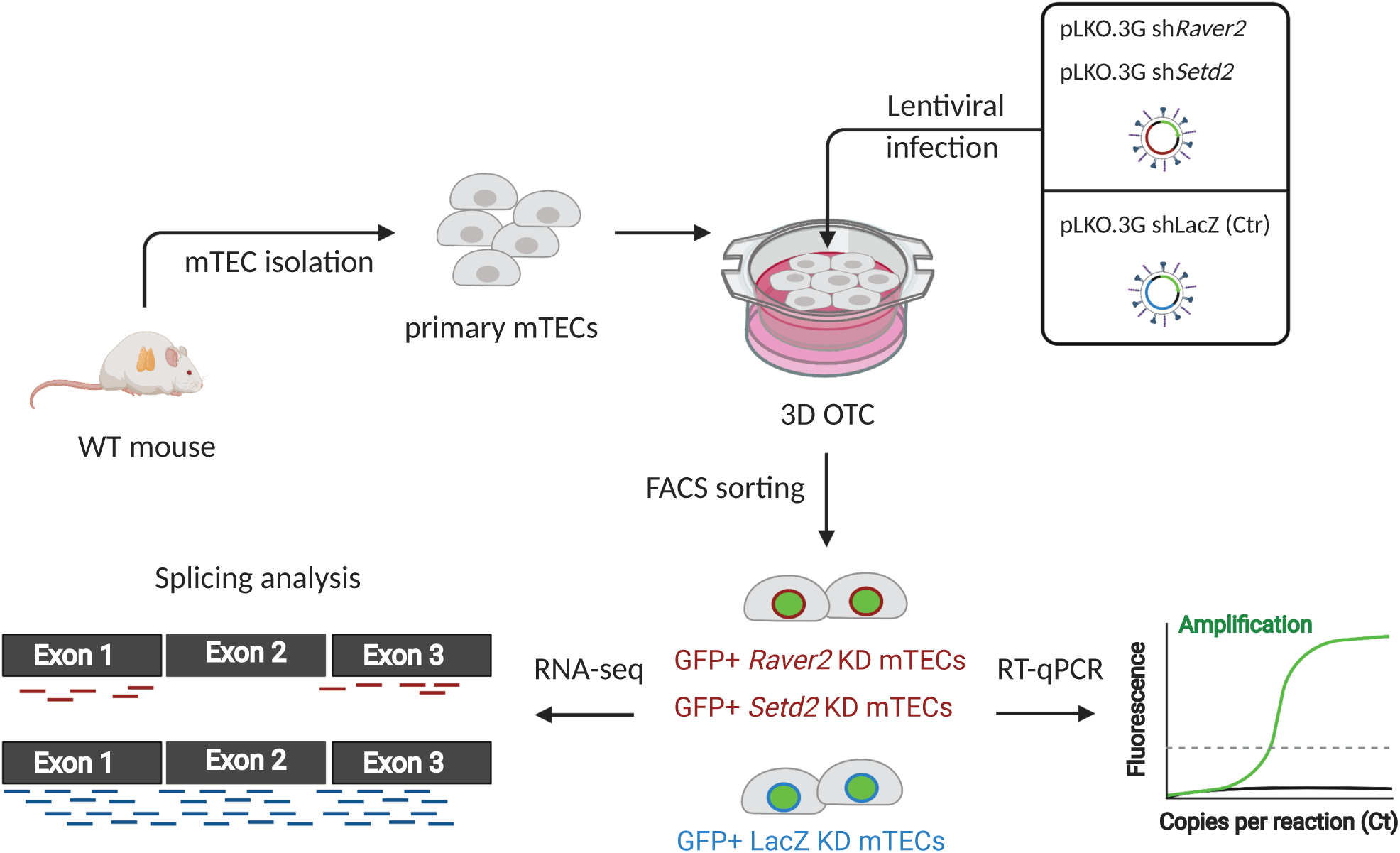
Schematic of the knockdown strategy of primary mTECs ex-vivo. Primary mTECs were isolated from thymi of WT mice and seeded along with shRNA-containing lentiviruses on a 3D organotypic culture system (3D OTC). Three days or five days after infection, GFP+ mTECs expressing the lentigenes were isolated for knockdown efficiency by qPCR or for splicing analyses by RNA-seq experiments. This figure was created with BioRender (biorender.com).

**Figure Sup 7.**
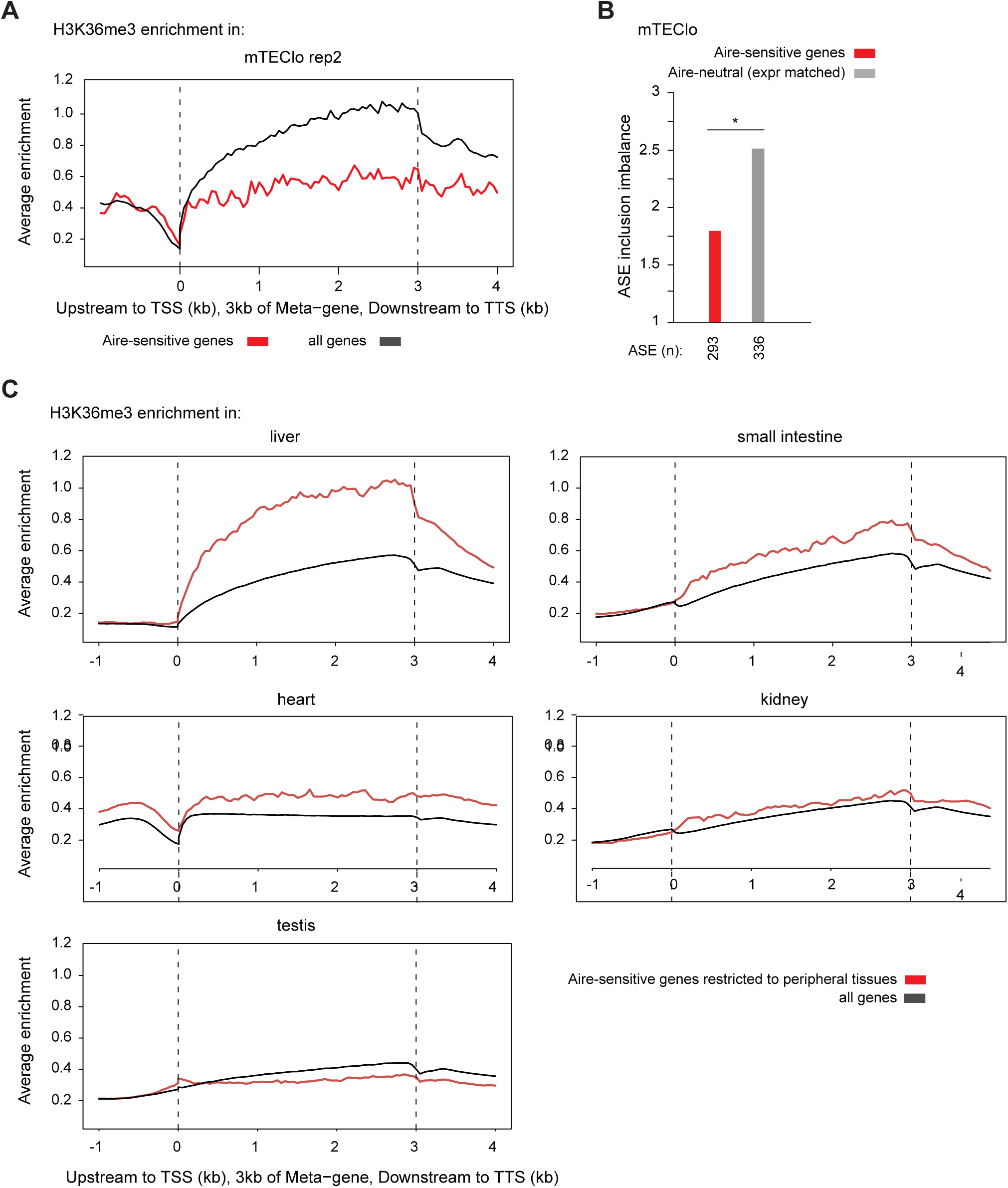
Reduced H3K36me3 deposition at Aire-sensitive genes in mTEClo in comparison to their tissues of expression. Metagene profiles of the average normalized enrichment of H3K36me3 for Aire-sensitive genes (red) in mTEClo **(A)** and in their tissues of expression **(C)**; all genes are shown in (black). (B) Levels of ASE inclusion imbalance for Aire-sensitive genes and neutral genes (expression matched), *P< 0.05 (Chi-squared test).

